# Positive feedback between demographic and selective fluctuations can greatly amplify the random fluctuation of population size

**DOI:** 10.1101/2024.06.13.598821

**Authors:** Yuseob Kim

**Affiliations:** Division of EcoScience, Ewha Womans University, Seoul, Korea 03760

**Keywords:** fluctuation, storage effect, population size, positive feedback, eco-evolutionary

## Abstract

Population sizes fluctuate over time probably due to random variability in the external environment. However, the severity of such fluctuations should depend on the characteristics of a species shaped in its evolutionary history. Previous studies have suggested that species are likely to evolve to minimize demographic fluctuations because an allele causing a smaller variance in offspring number is advantageous. However, this study finds that evolution in the opposite direction, favoring a mutation causing larger fitness fluctuation, occurs in a simple eco-evolutionary model under a randomly changing environment. This requires that (1) the mutant allele is under fluctuating selection within a subset, the field, of the population but neutral in another subset, the refuge, and (2) the field-to-refuge ratio of the carrying capacity is positively correlated to the fitness of the mutant allele. A general condition for the fixation of such a mutation was derived to depend on the relative strengths of demographic and fitness fluctuations and the mutational effect on the carrying capacity. Multi-locus simulations revealed that positive feedback between demography and selection accelerates the sequential fixations of fluctuation-amplifying mutations, leading to a drastic size fluctuation. This study therefore offers an unconventional explanation for large demographic fluctuations.

## INTRODUCTION

Natural populations experience temporal changes in abiotic and biotic environments. A randomly variable environment is expected to produce fluctuations in various aspects of demography, most importantly the number of reproducing individuals (i.e., population size), as well as fluctuation in the fitness effects of non-neutral or phenotype-changing alleles segregating in the population. If fitness differences under fluctuation are large, evolutionary changes may occur rapidly at the same speed as short-term ecological/demographic fluctuations. Recent developments in evolutionary biology have revealed that such rapid evolutionary changes are widespread (Bell 2010; Thompson 2013; Messer *et al*. 2016). One may then ask whether mutual interaction between demographic and selective fluctuations creates qualitatively novel eco-evolutionary dynamics that cannot arise if one of the demographic or population genetic variables is held constant.

Considering that a fluctuation in population size reflects a fluctuation in the reproductive success of individuals and that long-term reproductive success is maximized if its variance in the offspring number over time is minimized (Slatkin 1974; Gillespie 1977; Tuljapurkar 1982), one may predict that a population will evolve to reduce size fluctuation as much as possible (Pfister 1998). For example, a variant increasing the size rather than the number of eggs when plentiful resources are available may be positively selected as the offspring can survive better during an unfavorable period. Then, when a large fluctuation in population density, such as mast seeding or insect outbreak, does occur despite the advantage of a small temporal variance in offspring number, this may be seen as a non-adaptive response of the population to a heavy fluctuation of the biotic and/or abiotic environment (Myers 1988; Dwyer *et al*. 2004; White 2008; Bogdziewicz *et al*. 2020).

However, recent studies suggested that a population may evolve toward a larger fluctuation in its size. Liu *et al*. (2019) found that a reproductive strategy increasing the temporal variance of offspring number can be advantageous in populations reproducing in overlapping generations under various patterns of environmental fluctuations. Kim (2023) showed that a mutation amplifying the cyclic fluctuation of population size can be positively selected if this population is partially protected from fluctuating selection (i.e. fitness fluctuation occurs only in a subset of the population) at a given time. An additional condition promoting positive selection in the latter study was that the oscillation of fitness is positively correlated with that of population size and the regulation of population density is not strong enough so that the population size can over– or undershoot the carrying capacity. It is argued that the condition for partial protection from fluctuating selection is commonly satisfied in many species. The amplitude of fluctuation in selective pressure is likely to be heterogeneous over the geographic range of a spatially structured species. In plants and animals that have their life cycles divided into several stages and reproduce in overlapping generations, it is difficult to imagine a selective pressure uniformly acting on all individuals in the population regardless of the stage or age. For example, if a trait under fluctuating selection is expressed in the insect larval stage, alleles affecting this trait currently carried by adults are not visible to the selection. Similarly, alleles affecting traits that appear after germination remain neutral in the seed bank of a plant species. Such a subset of the population protected from fluctuating selection may be called a “refuge.” The presence of a refuge of various kinds was found to generate the advantage of rare alleles, a form of balancing selection known as the storage effect (Chesson & Warner 1981; Ellner & Hairston 1994; Reinhold 2000; Gulisija & Kim 2015; Gulisija *et al*. 2016; Bertram & Masel 2019; Yamamichi *et al*. 2023). However, Kim (2023) showed that if the population size is oscillating the negative frequency-dependent selection may not persist but turn into directional selection so that polymorphism is no longer maintained, unless fitness fluctuation is strong relative to population size fluctuation.

The theoretical results above — under what conditions balancing or directional selection on the alleles of fluctuating fitness will occur — were obtained from a model assuming cyclical/seasonal fluctuation in the environment and no reproduction in the refuge. A model with seasonal fluctuation is suitable for cases in which an environmental (seasonal) cycle spans multiple generations. However, for many species, one generation is not shorter than a seasonal cycle. Consequently, this study built a simple mathematical model in which the environment fluctuates randomly over time, which is probably relevant to a wider range of species including plants that exhibit mast seeding. In addition, the demographic assumption (no reproduction) of the refuge in Kim (2023), which was for specifically modeling a seed bank (Turelli *et al*. 2001), was relaxed so that it may represent a separate subpopulation undergoing reproduction and fluctuation in its carrying capacity. A general condition for balancing versus directional selection was analytically derived for the one-locus evolutionary dynamics under random environmental fluctuation. This one-locus model was then extended to a multi-locus model. The simulation of the latter showed that mutant alleles causing wider fluctuation in offspring number can reach fixation at multiple loci in a wide range of conditions. These fixations were found to occur at faster rates with increasing numbers of loci, as fluctuating population size and fluctuating selection reinforced each other in positive feedback. This result shows that eco-evolutionary dynamics arising under the general condition of heterogeneous fluctuating selection can produce highly variable demography.

## METHODS

### Eco-evolutionary model

Consider a haploid population that reproduces in discrete generations. It is divided into two subpopulations, termed the field and the refuge. A fluctuating environment causes fluctuations in the carrying capacities of both the field and refuge, which are given by 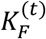 and 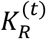 respectively for generation *t*. It is assumed that the carrying capacities of both subpopulations depend on common environmental factors that fluctuate randomly over time but that subpopulations may differ in the strengths of the dependence. The sizes of subpopulations, 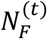 and 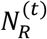, after the reproduction step in generation *t*, may or many not match the corresponding carrying capacities depending on the strength of population density regulation (see below). Before a new generation starts, each individual in both subpopulations is assumed to migrate to or remain in the field with probability 1 - *r* and migrate to or remain in the refuge with probability *r*. As a result, the proportion of the refuge in the total population is always *r* at the start of a generation. Next, the size of the field (refuge) changes from 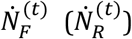 at the beginning to 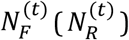 at the end of generation *t*, where 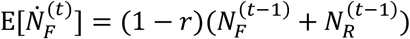 and 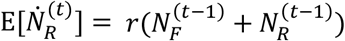.

It is assumed that the population is initially fixed for the *A*_1_ allele at a locus. Then, a new mutant allele *A*_2_, subject to fluctuating selection in the field but neutral in the refuge, arises. The fitness of an individual carrying *A*_2_ relative to *A*_1_ is given by 1 + *S*_*t*_ in the field, where E[*S*_*t*_] = 0 and Var[*S*_*t*_] > 0. Notably, while the arithmetic mean of the fitness of *A*_2_ over time is E[1 + *S*_*t*_] = 1, its geometric mean is less than 1. Therefore, natural selection is expected to eliminate this mutant allele from a population in a simple demographic model (containg the field only).

Let *n*_1F_ and *n*_2F_ be the numbers of individuals in the field carrying *A*_1_ and *A*_2_ alleles before reproduction for generation *t* (thus 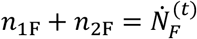). In the reproduction step, two modes of natural selection (soft vs. hard selection; Wallace (1975)) are considered regarding the effects of mutations on the population size. First, under soft selection, very strong density regulation is assumed so that the final size of the field is constrained to 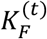. Each parent produces a Poisson number of progenies with mean 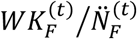, where *W* is 1 for *A*_1_ and 1 + *S*_*t*_ for *A*_2_, and 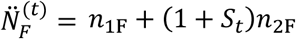. (Reproduction in Poisson numbers was needed to accommodate genetic drift; see below.) Second, under hard selection, fitness-changing mutations lead to changes in not only allele frequencies but also in the population size; they may increase or decrease the absolute reproductive output of the carriers such that the population size can increase over or decrease below the carrying capacity, which is now interpreted as the expeced population size assuming that all individuals carry the *A*_1_ allele. To this effect, each parent is now assumed to produce a Poisson number of progenies with mean 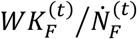. Namely, when *q* = *n*_2F_ /(*n*_1F_ + *n*_2F_) is the relative frequency of *A*_2_ at the beginning of a generation, the size of the field is expected to reach 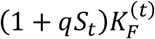. Reproduction in the refuge occurs similarly but without selection: each individual produces a Poisson offspring number with mean 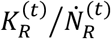.

Finally, how the environment fluctuates over time and how demographic and selective parameters 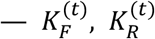, and *S*_*t*_ — respond need to be specified. The environmental condition is assumed to fluctuate randomly in each generation. More specifically, fluctuations occur such that 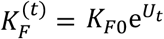 and 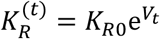 where *U*_*t*_ and *V*_*t*_ are drawn each generation from a joint normal distribution satisfying E[*U*_*t*_] = E[*V*_*t*_] = 0 and Cov[*U*_*t*_, *V*_*t*_] ≥ 0. The oscillation of carrying capacities is therefore symmetrical in the log scale, which is generally predicted in demographic models and observed in nature (Tuljapurkar & Orzack 1980; Myers 1988). *S*_*t*_ fluctuates such that E[*S*_*t*_] = 0, Cov[*S*_*t*_, *U*_*t*_] ≥ 0, and Cov[*S*_*t*_, *V*_*t*_] ≥ 0. Assuming non-zero correlations among the three parameters means that the population sizes and fitness respond to the fluctuation of common environmental variables.

### Stochastic simulation

Stochastic simulations of the above single-locus eco-evolutionary model and its extension to the multi-locus model were implemented in Mathematica ver. 11.3 (notebook files available at DOI 10.5281/zenodo.125116522). An individual has *L* loci that are arranged linearly with probability *c* for recombination between adjacent loci per generation. Simulation keeps track of two sets of lists: one containing the sets of haplotypes carried by non-zero individuals in the field and in the refuge (***H***_F_ and ***H***_R_) and the other containing the corresponding numbers of individiuals (***N***_F_ and ***N***_R_). Generation *t* begins after the migration step (re-distribution of individuals produced in the previous generation into the field and refuge) is completed. Haplotype numbers are first updated by mutation and recombination. A Poisson variate with mean *NL*μ, where *N* is the current size of a subpopulation at hand, is drawn as the total number of mutation events and each event is assigned to haplotypes and loci proportional to their current frequencies. Similarly, for *L* > 1, a Poisson variate with mean *N*(*L*-1)*c* is drawn as the total number of recombination events in the subpopulation. For each recombination event, two haplotypes are chosen proportional to their frequencies and one of the *L*-1 intervals is chosen to be the position of crossing-over between the two haplotypes. Newly created or lost haplotypes during mutation and recombination steps are tracked by updating (***H***_F_, ***H***_R_) and (***N***_F_, ***N***_R_). Then, the updated lists are further updated in the step of reproduction according to their fitness. Under soft selection, the number of individuals with haplotype *i* born in the next generation is Poisson distributed with mean 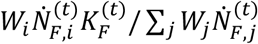 in the field and 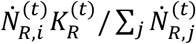 in the refuge, where *W*_*i*_ is the relative fitness of haplotype *i*. Under hard selection, the corresponding mean number in the field is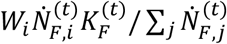. Finally, when there are *N*_F_ and *N*_R_ individuals in the field and refuge, Poisson variates with means *rN*_F_ and (1-*r*)*N*_R_ are drawn to determine the numbers of migrants from field to refuge and from refuge to field, respectively. Migrants are chosen proportional to the haplotype frequencies and then merged with the residents of the destination. This completes a single-generation iteration in updating haplotype lists.

## RESULTS

### Analytic results: the fate of the *A*_2_ allele

This section aims to predict the fate of the rare *A*_2_ allele — whether it will be lost, fixed, or increased to intermediate frequencies and maintained polymorphic in the population — as a function of the demographic and fitness parameters. First, soft selection with strong density regulation, under which 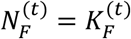 and 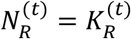, and no genetic drift are assumed. Let *n*_1_ and *n*_2_ be the total numbers of individuals (counted after reproduction/selection and before migration) carrying *A*_1_ and *A*_2_ in generation *t* – 1 and *n’*_1_ and *n’*_2_ be their expected numbers in generation *t*. Then, with *n*_1_ ≫ *n*_2_,

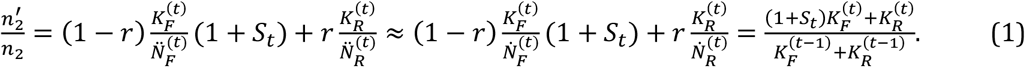

This equation takes it into account that the change in the absolute number of *A*_2_ is determined first by demography, as the size of field changes from 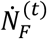 to 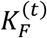 and that of refuge from 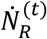 to 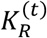, and also by fitness 1 + *S*_*t*_. Whether the copy number of rare allele *A*_2_ is expected to increase or decrease is determined by the expectation of log 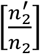 over demographic and fitness fluctuations. It is shown in Appendix A that, the condition for the *A*_2_ allele being positively selected (i.e. invading the population initially fixed for *A*_1_), 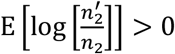, is simplified to

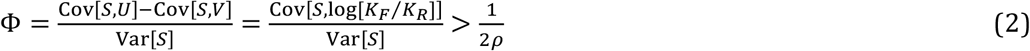

where *ρ* = *K*_*R*0_/*K*_*F*0_ (note that 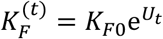 and 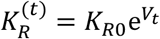) represents the relative size of the refuge. Therefore, the rare *A*_2_ allele is positively selected if the covariance between its fitness and the log-ratio of the field to refuge size is large relative to the amplitude of the fitness fluctuation. This requires that the demographic fluctuation of the field be sufficiently large, as already shown in Kim (2023), and that of the refuge is either small or out of phase with the fluctuation of the field. This condition is easier to meet with the increasing size of the refuge.

In a similar manner, the condition for the *A*_1_ allele to invade a population initially fixed for *A*_2_ under soft selection becomes

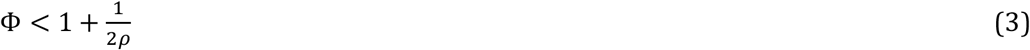

(Appendix A). From inequalities (2) and (3), it is predicted that the *A*_1_ and *A*_2_ alleles coexist in the population if 1/(2*ρ*) < Φ < 1 + 1/(2*ρ*) under soft selection. With Φ < 1/(2*ρ*), the *A*_2_ allele is eliminated from the population, as the fitness fluctuation that reduces its geometric mean in the field is more important than population size fluctuation. With Φ > 1 + 1/(2*ρ*), the *A*_2_ allele is predicted to reach fixation in the population.

Using the same approach, the fate of the *A*_2_ allele under hard selection can be analyzed. Note that the carrying capacity of the field is given by 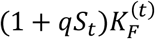 with hard selection where *q* is the frequency of *A*_2_. The frequency change of *A*_2_ when rare is therefore effectively identical to that in soft selection. Therefore, the condition for *A*_2_ invading the population fixed for *A*_1_ is again Φ > 1/(2*ρ*). However, the condition for the fixation of *A*_2_ is drastically different from that under soft selection. It is shown in Appendix A that the condition for 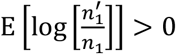 for rare *A*_1_ is exactly the condition for 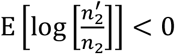 for rare *A*_2_. Therefore, the condition for the *A*_1_ allele to invade the population fixed for *A*_2_ is approximately Φ < 1/(2*ρ*), which means that the fixation of *A*_2_ is ensured if Φ > 1/(2*ρ*): once positively selected after its appearance by mutation, the *A*_2_ allele is expected to increase in frequency all the way to fixation. This leads to a fluctuation amplification in the size of the field, from 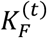 to 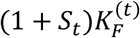. This result is consistent with the finding in Kim (2023) that, under weak density regulation of the population size, an allele with a wider oscillation in absolute fitness reaches fixation in the presence of refuge.

### One-locus simulation

To validate and explore the dynamics of the above eco-evolutionary model, the one-locus stochastic simulation was performed. The demographic and fitness fluctuations were given by *U*_*t*_ = *a*_*U*_*Z*_1_ + *b*_*U*_*Z*_2_, *V*_*t*_ = *a*_*V*_*Z*_1_ + *b*_*V*_*Z*_3_, and *S*_*t*_ = *a*_*S*_*Z*_1_ + *b*_*S*_*Z*_4_, where the *Z*_*i*_s are standard normal variates that were drawn independently each generation. Therefore, population sizes and fitness were modeled to respond to the fluctuation of a common environmental determinant represented by *Z*_1_. In this setting, 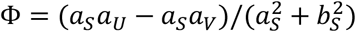.

Simulations were first performed with soft selection. Initially the numbers of *A*_1_ and *A*_2_ individuals changed deterministically once the demographic and fitness parameters (*U, V*, and *S*) for a given generation were sampled from their joint distribution, effectively assuming that the sizes of the field and refuge were finite and fluctuating but very large so that genetic drift can be ignored (Fig. 1A). Various values of *a*_*S*_ and *b*_*S*_ were chosen so that Φ ranged from 0.3 to 2, while ρ was set to 1. Starting from 0.5, the frequency of the *A*_2_ allele at the 5,000th generation was recorded for 1000 replicates per parameter set. As predicted, with Φ ≤ 0.5, the frequency of *A*_2_ approached 0, thus confirming negative selection, and with Φ ≥1.5 the frequency approached 1, thus confirming positive directional selection. The distribution of the allele frequency was approximately uniform between 0 and 1 when Φ = 1.

**Figure 1.**
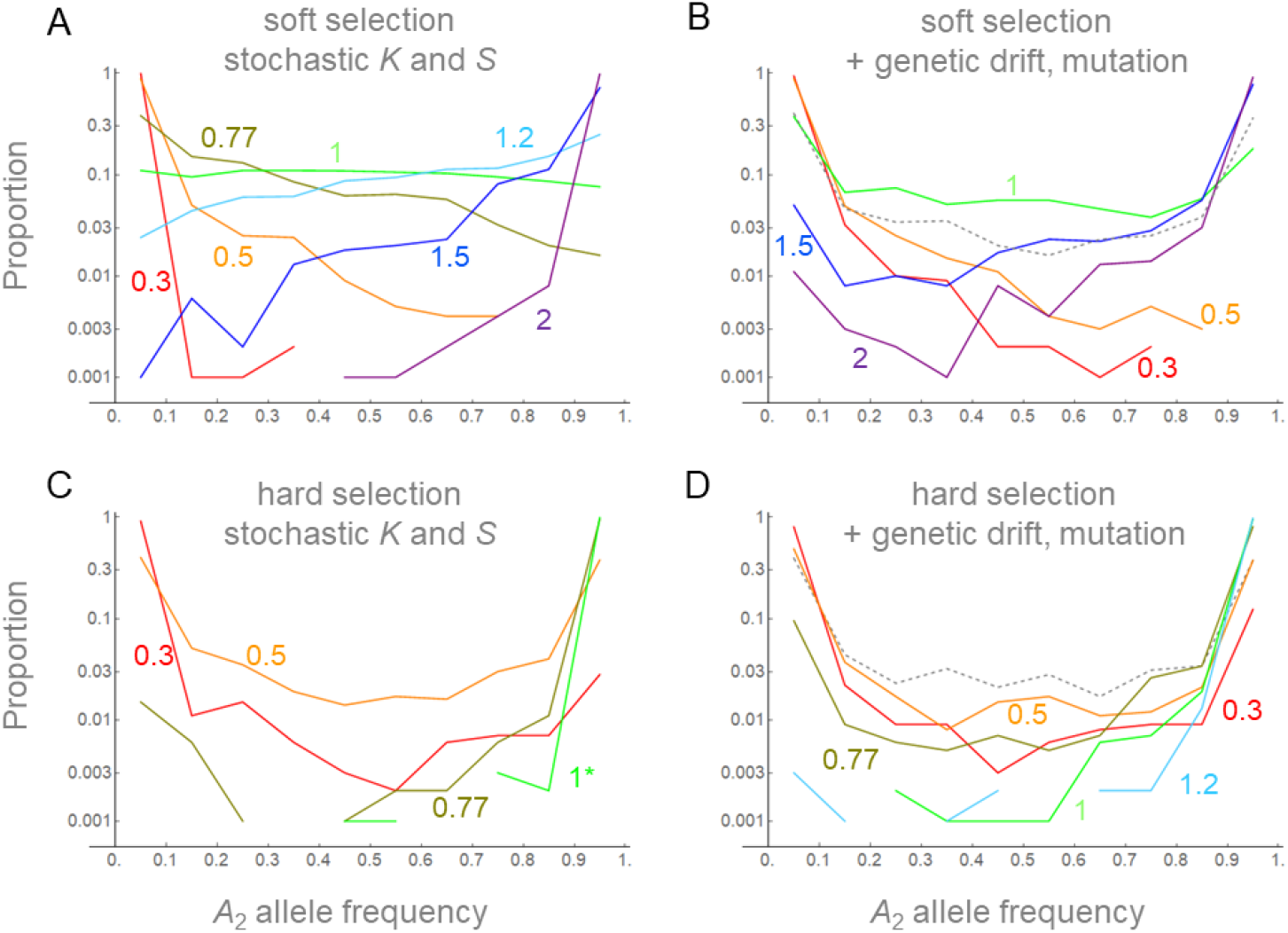
Distribution of *A*_2_ allele frequencies (*p*) at the 5,000^th^ generation in the stochastic simulations of soft (A, B) and hard (C, D) selection. The initial frequency of *A*_2_ is 0.5 in both field and refuge. Reproduction occurred while 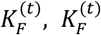 and *S*_*t*_ varied stochastically without (A, C) or with (B, D) genetic drift and bidirectional mutation (μ = 5 × 10^−5^). Proportions of simulation replicates (out of 1000) with the frequencies in the interval 0 ≤ *p* < 0.1, 0.1 ≤ *p* < 0.2, …, 0.8 ≤ *p* < 0.9, and 0.9 ≤ *p* ≤ 1 were plotted and connected by lines. Fluctuations of population sizes are given by *K*_*R*0_ = *K*_*F*0_ = 1000 (*ρ* = 1), *a*_*U*_ = 0.3, *b*_*U*_ = 0.1, *a*_*V*_ = 0.05, *b*_*V*_ = 0.1. Parameters of fluctuation were chosen to yield Φ = 0.3, 0.5, 0.77, 1, 1.2, 1.5, and 2: (*a*_*S*_, *b*_*S*_) = (0.07, 0.23), (0.1, 0.2), (0.14, 0.16), (0.2, 0.1), (0.2, 0.04), (0.15, 0.05), and (0.1, 0.05). In the case of hard selection with Φ = 1 and no genetic drift (panel C), simulation ran only upto 2,500 generations: longer simulations or larger Φ resulted in all replicates with final frequencies > 0.9.

Next, genetic drift and bidirectional mutation between two alleles were added to the above simulations (Fig. 1B). This weakened the strengths of all modes of selection. With Φ = 1, more simulation runs ended with allele frequencies close to 0 and 1. However, the proportion of runs ending with intermediate allele frequencies was still larger than that observed when the alleles were set as neutral (dashed curve in Fig. 1B). Therefore, an evolutionary force to maintain polymorphism beyond the level under the neutrality, namely balancing selection, was confirmed.

Similarly, the fate of the *A*_2_ allele under hard selection was examined. Stochastic simulation confirmed the approximation above predicting that *A*_2_ is positively selected towards fixation if 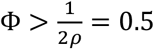, although genetic drift and mutation weakened the trend (Fig 1C,D). Positive selection on both rare and common *A*_2_ under the same condition means that this allele may not remain polymorphic for long in the population. Indeed, across all values of Φ there were fewer simulation runs ending with intermediate frequencies of *A*_2_ compared to the simulation of neutral alleles subject to genetic drift and bidirectional mutations only (dashed curve in Fig. 1D). Therefore, balancing selection that was observed with soft selection did not arise with hard selection.

### Multi-locus simulation

To investigate how the eco-evolutionary dynamics shown in the single-locus model above play out when there are many such loci in the genome, new sets of simulation were performed in which a haploid individual is modeled to have *L* linked loci, each carrying either the *A*_1_ or *A*_2_ allele described above. The simulation tracks the numbers of individuals of different haplotypes (see METHODS). Epistasis is not considered; the relative fitness of haplotype *i* is given by 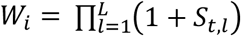, where *S*_*t,l*_ is 0 if the *l*th locus of the haplotype carries *A*_1_ and *a*_*S*_*Z*_1_ + *b*_*S*_*Z*_4_ if *A*_2_. *Z*_1_ is drawn once for all loci while *Z*_4_ is drawn separately for different loci. Individuals randomly pair and undergo recombination by crossing-over such that the probability of recombination between adjacent loci is *c* per generation. The simulation starts with a population fixed for *A*_1_ at all loci. Then *A*_2_ arises by mutations with probability μ/locus/generation.

Simulations can be run under two different parameter domains. First, the demographic and selective parameters that lead to balancing selection in the one-locus simulation (soft selection with 1/(2*ρ*) < Φ < 1 + 1/(2*ρ*)) were chosen. In limited sets of simulation (*L* = 4, or 10), the observed level of per-locus polymorphism was above the level in equivalent simulations with neutral alleles (*S*_*t*_ = 0), thus confirming the operation of multi-locus balancing selection. This result is described in Appendix B. This study does not further explore this but focuses on the second (and wider) parameter domain, namely Φ > 1 + 1/(2*ρ*) with soft selection and Φ > 1/(2*ρ*) with hard selection, in which the fixation of the *A*_2_ allele was observed in the single-locus simulation. As expected, substitutions occurred sequentially until *A*_2_ reached fixation at all *L* loci (Fig. 2 and Fig. S1 for example). Times (in generations) when the frequency of *A*_2_ first exceeded 0.9 at each locus were recorded and sorted into *T*_1_ to *T*_*L*_. There is a large difference in substitution dynamics between soft and hard selection. With soft selection, the mean waiting time, 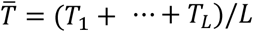, increased moderately as *L* increased from 1 to 8 (Fig. 3A). This may be explained by a reduction in the efficacy of positive selection on *A*_2_ at a given locus when selection at other loci results in genome-wide reduction in the effective population size.

**Figure 2.**
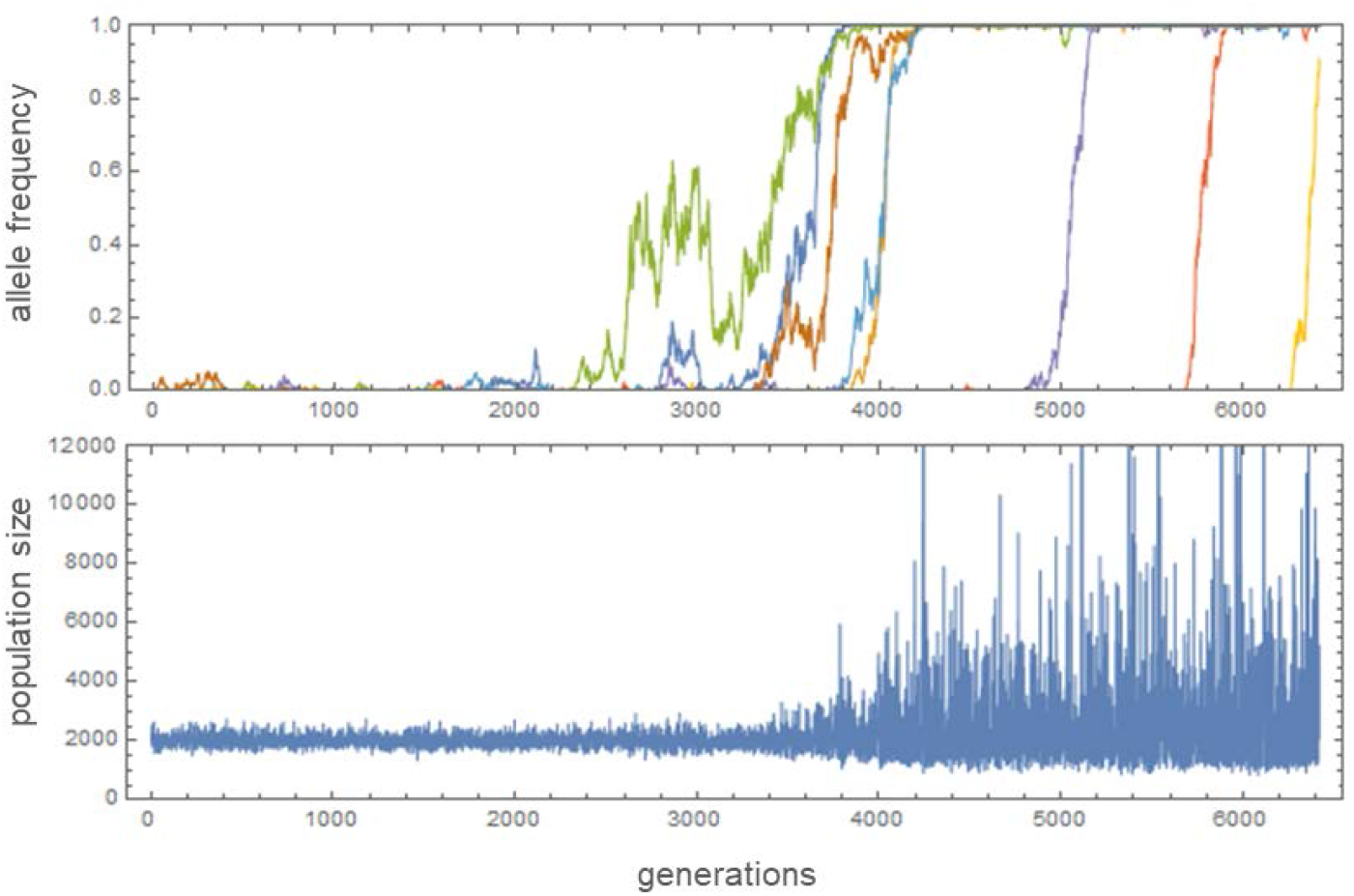
Changes of mutant (*A*_2_) allele frequencies and population size 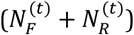 in a multi-locus simulation of hard selection with *L* = 8, *c* = 0.02, *K*_*R*0_ = *K*_*F*0_ = 1000, μ = 2 ×10^−5^, and Φ = 1 (*a*_*S*_ = 0.15, *b*_*S*_ = 0, *a*_*U*_ = 0.15, *a*_*V*_ = 0, *b*_*U*_ =0, *b*_*V*_ = 0.1). Allele frequency trajectories of 8 loci were plotted in different colors.

**Figure 3.**
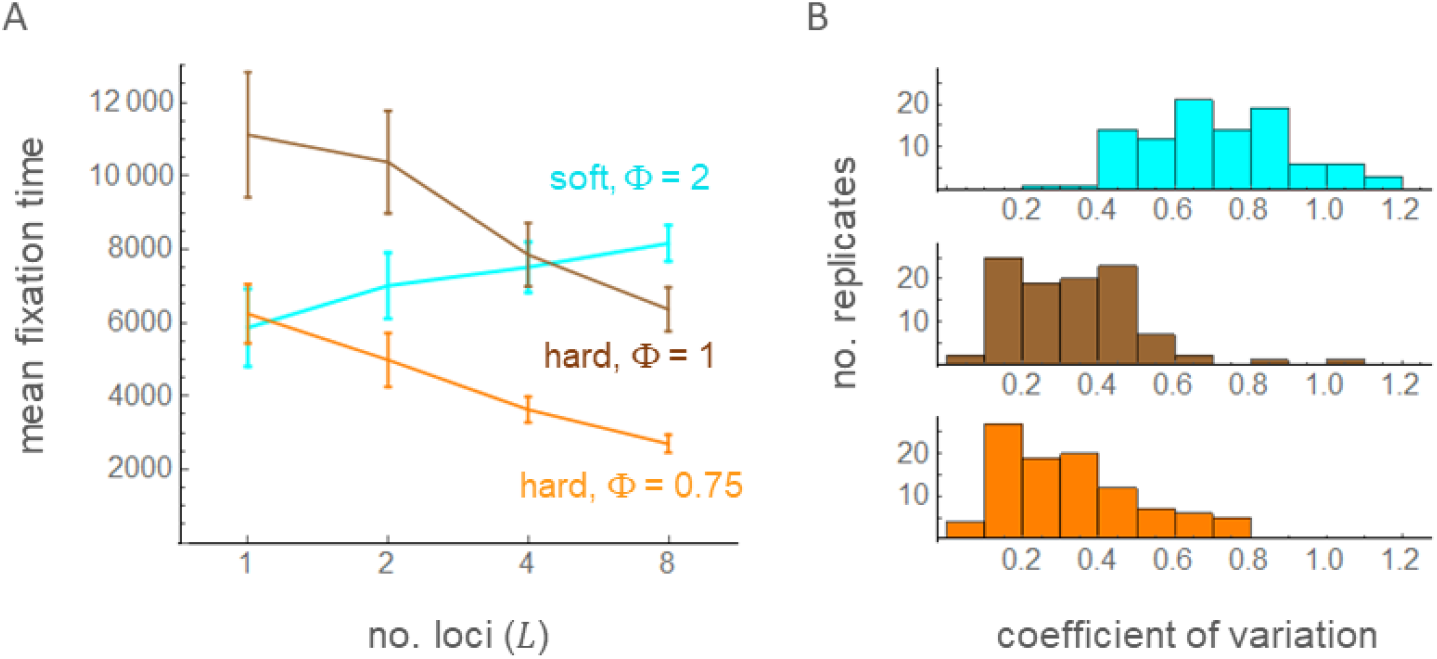
Mean fixation time 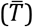 with varying number of loci (A; *L* = 1, 2, 4, and 8) and the coefficient of variation in fixation times with *L* = 8 in multi-locus simulation. Error bars show two times the standard errors. A total of 100 replicates of simulations were run under soft selection with Φ = 2 (cyan; *a*_*S*_ = 0.1, *b*_*S*_ = 0, *a*_*U*_ = 0.25, *b*_*U*_ = *a*_*V*_ = *b*_*V*_ = 0.05), hard selection with Φ = 1 (brown; *a*_*S*_ = 0.1, *b*_*S*_ = 0, *a*_*U*_ = 0.1, *a*_*V*_ = 0, *b*_*U*_ = *b*_*V*_ = 0.1), or hard selection with Φ = 0.75 (orange; *a*_*S*_ = 0.2, *b*_*S*_ = 0, *a*_*U*_ = 0.25, *a*_*V*_ = 0.1, *b*_*U*_ = *b*_*V*_ = 0.05). Other parameters are identical to those in Figure 3.

With hard selection, however, 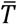 decreased as *L* increased. Furthermore, the coefficient of variation, the standard deviation of *T*_*i*_’s divided by 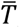, was significantly smaller with hard selection than with soft selection (Fig. 3B). As expected for hard selection, the amplitude of the population size fluctuation progressively increased as the number of loci fixed for *A*_2_ increased (Fig. 2). These results clearly demonstrate that positive feedback between demographic and selective fluctuations under hard selection led to a great amplification of population size fluctuation; an initial fluctuation in 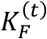 positively selected the mutant allele *A*_2_ at one locus whose fitness fluctuates in positive correlation with 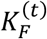, leading to the fixation of *A*_2_ and therefore a larger fluctuation of the field (given by 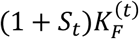, which makes it easier for the *A*_2_ allele at another locus to be positively selected. This process could continue to increase the fluctuation of the field until its carrying capacity is given by 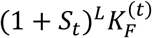, where *L* is the number of loci available in the genome for mutation into *A*_2_. There was also a moderate effect of the recombination rate in accelerating the multi-locus fixation of *A*_2_ with hard selection (Table S1). Fixation times generally increased with decreasing recombination rates for both soft and hard selection, suggesting that tighter linkage increased the effect of clonal interference.

## DISCUSSION

A key assumption in this study is that a tradeoff commonly occurs for the fitness effects of a mutant allele between different time points in the environmental fluctuation. Namely, a mutant phenotype that increases the reproductive output relative to others when the environment is generally favorable is likely to decrease it relative to others during an unfavorable period. Such a mutant allele increases the variance of the offspring number over time while the mean may remain unchanged. For example, as postulated by Slatkin (1974), a mutation that makes a female bird lay 10 eggs in one batch instead of spreading them over time in a randomly and rapidly fluctuating environment will increase the variance of her fitness while the mean remains the same. Modeling the fluctuating fitness of a mutant by 1 + *S*_*t*_ reflects such a tradeoff. Classical theory predicts that such a mutant (termed an arithmetic-mean-preserving or AMF allele in Kim (2023)) is eliminated from a population as its geometric mean fitness is less than 1. However, it was demonstrated in this study that the AMF mutant is positively selected if a correlation between demographic and fitness fluctuations exist in the presence of refuge. Analysis given in Appendix A shows that the arithmetic mean fitness of an AMF allele is effectively increased above 1 due to correlated demographic fluctuation in the field. However, without a refuge, its geometric mean cannot exceed 1. Namely, as pointed out in Kim (2023), by dampening the fluctuation of the AMF allele’s effective fitness in the total population the refuge turns the heightened arithmetic mean into the geometric mean greater than 1.

One may also model the fitness of a mutant by exp[*S*_*t*_], making the mutant (termed a geometric-mean-preserving or GMF allele) quasi-neutral as its geometric mean fitness is identical to that of the wild-type allele even without demographic fluctuation. Analyzing the model of cyclic environmental oscillation, Kim (2023) showed that the parameter range of positive selection for a GMF allele is wider than that for an AMF allele. Therefore, the major results obtained in this study, both balancing and positive directional selection on a fitness-amplifying allele, are probably easier to achieve with GMF mutations. However, a GMF allele may be considered simply a beneficial allele as the arithmetic mean of offspring number is greater than the wild-type allele. Such a mutation is probably rare and was not considered in this study.

The refuge in the current model may correspond to various demographic elements. It directly models a patch in a spatially structured population that is protected from fluctuating selection. However, given that many studies modeling the storage effects of a certain life-cycle stage yield equations similar to eq. (1), namely in the form *p*′/*p* = (1 − *r*)*f* + *r* where *p* is the frequency of the rare-type, *r* is the proportion of the refuge (e.g., adult stage when selection acts on the juveniles, seed bank, or diapausing eggs), and *f* is the relative fitness of the type outside the refuge (Chesson & Warner 1981; Ellner & Hairston 1994; Turelli *et al*. 2001; Gremer & Venable 2014), this study should apply to such species if 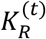 is given an appropriate interpretation. A major result in this study is that an AMF allele is positively selected if correlation of its fitness fluctuation with the size of the field is larger than that with the size of the refuge. This requirement is important for sex-limited gene expression. Previous studies suggested that if a gene is expressed in one sex the other sex could be a refuge in a dioecious species (Reinhold 2000; Kim 2023). However, as equal numbers of male and female gametes are used in reproduction, the population sizes of reproductive males and females are effectively equal while the sex ratio of the adults may fluctuate. Therefore, Φ is zero as the size of the field relative to the refuge does not fluctuate at all. Hence, positive selection on the fitness-amplifying mutation shown in this study should not happen for sex-limited genes, unless GMF mutants are considered.

While the eco-evolutionary model of this study can result in both balancing and directional selection on size-fluctuation-amplifying alleles, this study focused on the latter. The condition for the fixation of those alleles is less restrictive compared to that for balanced polymorphism, because it requires relatively weaker selection (smaller variance in relative fitness) for a given demographic fluctuation to satisfy Φ > 1 + 1/(2*ρ*) with soft selection and Φ > 1/(2*ρ*) with hard selection. Demographic fluctuation that drives this evolutionary process should apply equally to all loci in the genome. Therefore, fixations occur at all available loci on which proper mutations, i.e. the *A*_2_ allele, arise, as observed in the multi-locus simulation. With soft selection, positive selection at each locus is contributed by the same degree of demographic fluctuation regardless of the evolutionary changes at other loci. However, with hard selection, fluctuation in the size of the field affecting a given locus intensifies as more fluctuation-amplifying mutations reach fixation at other loci. Since the carrying capacity of the field is effectively 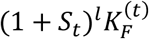, where *l* is the number of loci where *A*_2_ reached fixation, and wider fluctuation generates stronger positive selection on the rare *A*_2_ (larger E[Δ_2_] in Appendix A), fixation at one locus facilitates the subsequent fixations at other loci. This positive feedback between demographic fluctuation and positive selection drastically increases the amplitude of fluctuation in the total population size (Fig. 2). How much the amplitude will increase should depend on the number of loci at which mutations subject to hard selection arise. There should be multiple independent ways in which changes in mutants’ relative fitness can be translated into changes in the mean absolute fitness of the entire population. For various life history traits, one may imagine phenotypes (smaller eggs, smaller adult body, earlier onset of first reproduction, etc.) that increase the absolute number of surviving offspring during a favorable, resource-rich period but decrease the number in a bad period while not affecting the amount of resources that the carriers of the alternative, ancestral alleles acquire. Then, many loci may be available for hard selection, the combined effect of which may lead to a very large fluctuation in population size.

The size of a population and its fluctuation should be governed not only by extrinsic factors (the environmental variables such as temperature and parasite prevalence) but also by intrinsic factors (species-specific responses to those extrinsic factors). The intrinsic factors such as various life-history traits are the product of the species’ evolutionary history. For example, phenotypic plasticity responding to environmental cues can reduce the amplitude of the population size fluctuation, thus acting as a mechanism of population regulation (Caswell 1983; Reed *et al*. 2010), which according to the classical theory is adaptive as it enhances long-term reproductive success by increasing the geometric mean fitness (Gillespie 1977; Pfister 1998). If life history traits evolved in the direction toward suppressing random or cyclic fluctuations in the population size, fluctuations observed in nature should be interpreted as the effect of variable environments that the evolved mechanisms of population regulation were unable to contain. However, this study suggests that evolution can proceed in the opposite direction. A large random fluctuation in population size can be the product of adaptive evolution driven by positive feedback between demographic and selective fluctuations. When the degrees of population size fluctuation are different across species, the eco-evolutionary dynamics proposed in this study might be particularly important for those exhibiting large fluctuations. Interestingly, annual desert plants experiencing wider fitness fluctuations after germination have greater proportions of their population remaining in the seed bank (Gremer & Venable 2014). This observation is in agreement with the analytic results in this study because a larger seed bank corresponds to a larger value of ρ, which makes it easier for the fixation of the fluctuation-amplifying alleles (i.e., lowering the threshold of Φ for positive selection).

## ACKNOWLEDGMENT

This study was supported by the National Research Foundation (NRF) grant 2020R1A2C1009261 funded by the Korean government.

## Conflict of interest statement

The author declares no conflict of interest

## Appendix A. Derivation of conditions for balancing and directional selection on *A*_2_

Here only deterministic changes in allele frequencies are tracked as a very large population is assumed. At the end of reproduction but before migration into subpopulations at generation *t* - 1, there are *n*_1_ and *n*_2_ copies (individuals) of *A*_1_ and *A*_2_ in the entire population. After migration, the numbers of *A*_1_ and *A*_2_ copies are *n*_1*F*_ = (1 − *r*)*n*_1_ and *n*_2*F*_ = (1 − *r*)*n*_2_ in the field and *n*_1*R*_ = *rn*_1_ and *n*_2*R*_ = *rn*_2_ in the refuge. Therefore, the relative frequencies of alleles are identical between the field and refuge before reproduction at the beginning of generation *t*.

Soft selection is first considered. Then, each *A*_2_ allele produces 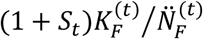 copies of descendants, where 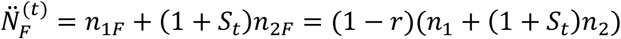. Since *n*_1_ » *n*_2_ after *A*_2_ appears by mutation, *n*_1_ + (1 + *S*_*t*_)*n* ≈ *n*_1_ and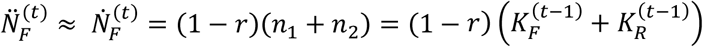. Similar approximations apply to the refuge. Let *n’*_2_ be the expected number of *A*_2_ in generation *t*. This leads to eq. (1). Then, with 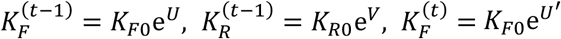, and 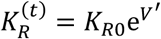,

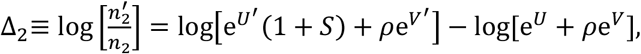

where *S* = *S*_*t*_ and *ρ* = *K*_*R*0_/*K*_*F*0_. If E[Δ_2_] > 0, where the expectation is over demographic and fitness fluctuations, the rare allele *A*_2_ is expected to increase its copy number over time. Applying the Taylor expansion up to the second order,

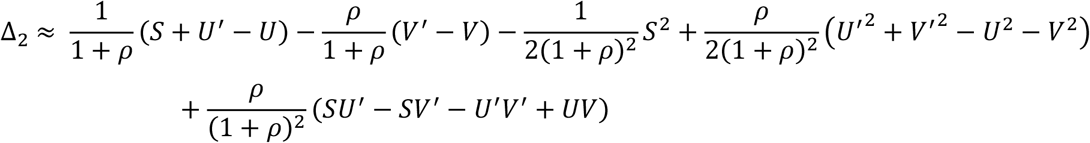

Since all random variables in the above equation have the mean at zero and (*U, V*) and (*U*^′^, *V*^′^) are from the same joint probability distribution,

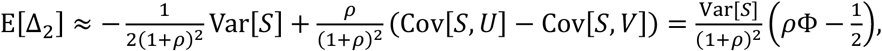

where

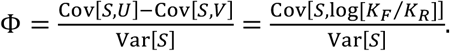

Then, the condition for the *A*_2_ allele being positively selected (i.e. invading the population initially fixed for *A*_1_), is E[Δ_2_] > 0 or 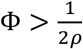.

A similar result can be obtained for the condition for the *A*_1_ allele to invade a population initially fixed for *A*_2_. From

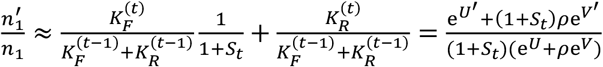

for *n*_1_ ≪ *n*_2_, the dynamics of the rare *A*_1_ allele depends on the expectation of

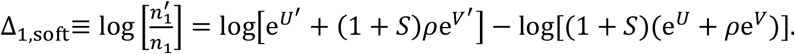

Then, using quadratic approximation

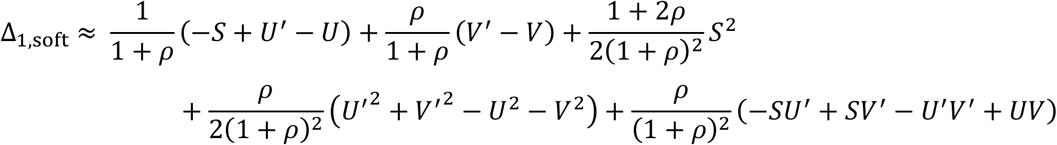

one obtains

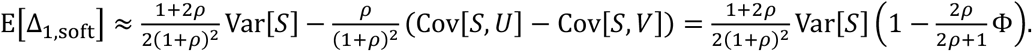

The condition for the rare *A*_1_ allele to increase in frequency is E[Δ_1,soft_] > 0 or

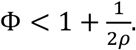

With hard selection, the frequency change of *A*_2_ when rare is again given by eq. (1). Therefore, the condition for *A*_2_ invading the population fixed for *A*_1_ (E[Δ_2_] > 0 above) is no different from that for soft selection. However, because the carrying capacity of the field fixed for *A*_2_ at generation *t* is given by 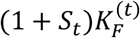 under hard selection, the opposite condition for *A*_1_ invading *A*_2_ is given by

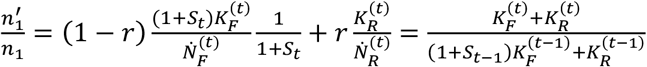

Then,

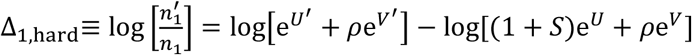

and, again, the quadratic approximation is obtained as

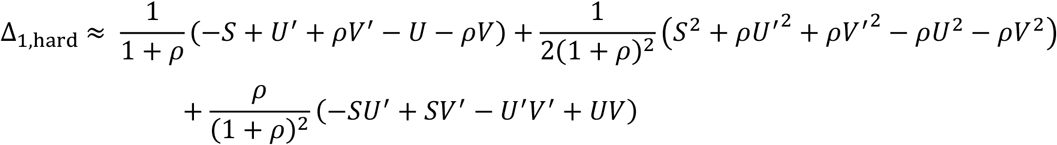

Finally,

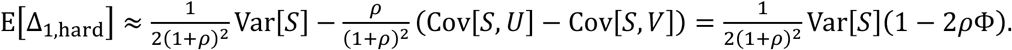

The condition for the fixation of *A*_2_ (the failure of *A*_1_ to invade) is E[Δ_1,hard_] < 0 or 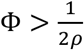.

Now, a heuristic argument can be given for positive selection (both soft and hard) on *A*_2_, which is expected to be eliminated from a population whose size does not fluctuate. Positive selection occurs because the mean number of offspring per parent, i.e. absolute fitness, of the mutant in the field is given by 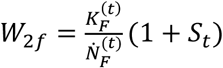 with fluctuation in population size (eq. 1). With positive correlation between 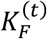 and *S*_*t*_, the fluctuation of *W*_2*f*_ becomes greater than that of 1 + *S*_*t*_ and, more importantly, its arithmetic mean can now exceed 1. However, the geometric mean of *W*_2 *f*_ is still not greater than 1 due to its fluctuation over time. The presence of a refuge then reduces the variance of mutant’s fitness averaged across the whole population such that the geometric mean of this average fitness becomes greater than 1. In summary, the refuge turns the heightened arithmetic mean raised by the demographic fluctuation of the field into an elevation of the geometric mean in the total population (see also Kim (2023))

## Appendix B. Multi-locus simulation for balanced polymorphism

Considering that (1) the negative frequency-dependent selection on a variant with fitness fluctuation was caused primarily by correlated fluctuation in the carrying capacity (population size) of the field and that (2) this demographic fluctuation should be “felt” at all loci in the genome, one may expect that such balancing selection can occur simultaneously at many loci harboring such variants. To verify if this is true, simulations were run under soft selection and demographic/selective parameters that promoted polymorphism in the single-locus model (Φ = 1 for ρ = 1) with *L* = 1, 4, or 10. Exemplary trajectories of allele frequencies are shown in Fig. S2. Each run lasted at least 10^5^ generations, over which the mean of expected heterozygosity (*H* = 2*p*(1 − *p*)) and the proportions of time *A*_2_ spent in frequency between 0.1 and 0.5, *P*_15_, and between 0.5 and 0.9, *P*_59_, were recorded per locus (Table S2). In all cases with fluctuating fitness (*a*_*S*_ > 0), polymorphism per locus was above the level observed in equivalent simulations with neutral alleles (*S*_*t*_ = 0), thus confirming multi-locus balancing selection. For cases of stronger selection (larger Var[*S*_*t*_]), the level of polymorphism per locus declined as *L* increased from 1 to 10. The combined effects of the number of loci and the rate of recombination were not easy to interpret as they depended on the parameters of selection (for sets of fitness/demographic parameters that yield the same Φ = 1, *H* responded differently with varying *L* and *c*). One possible explanation for generally lower variation with 10 loci instead of 1 or 4 is that the simultaneous segregation of alleles at many loci increased the variance of the offspring numbers, thus decreasing the effective population size and weakening the negative frequency-dependent selection that is needed for protected polymorphism. Further exploration of multi-locus polymorphism at a larger scale (in both number of loci and population size) may not be feasible with the current simulation approach based on haplotype frequencies.

**Table S1.**
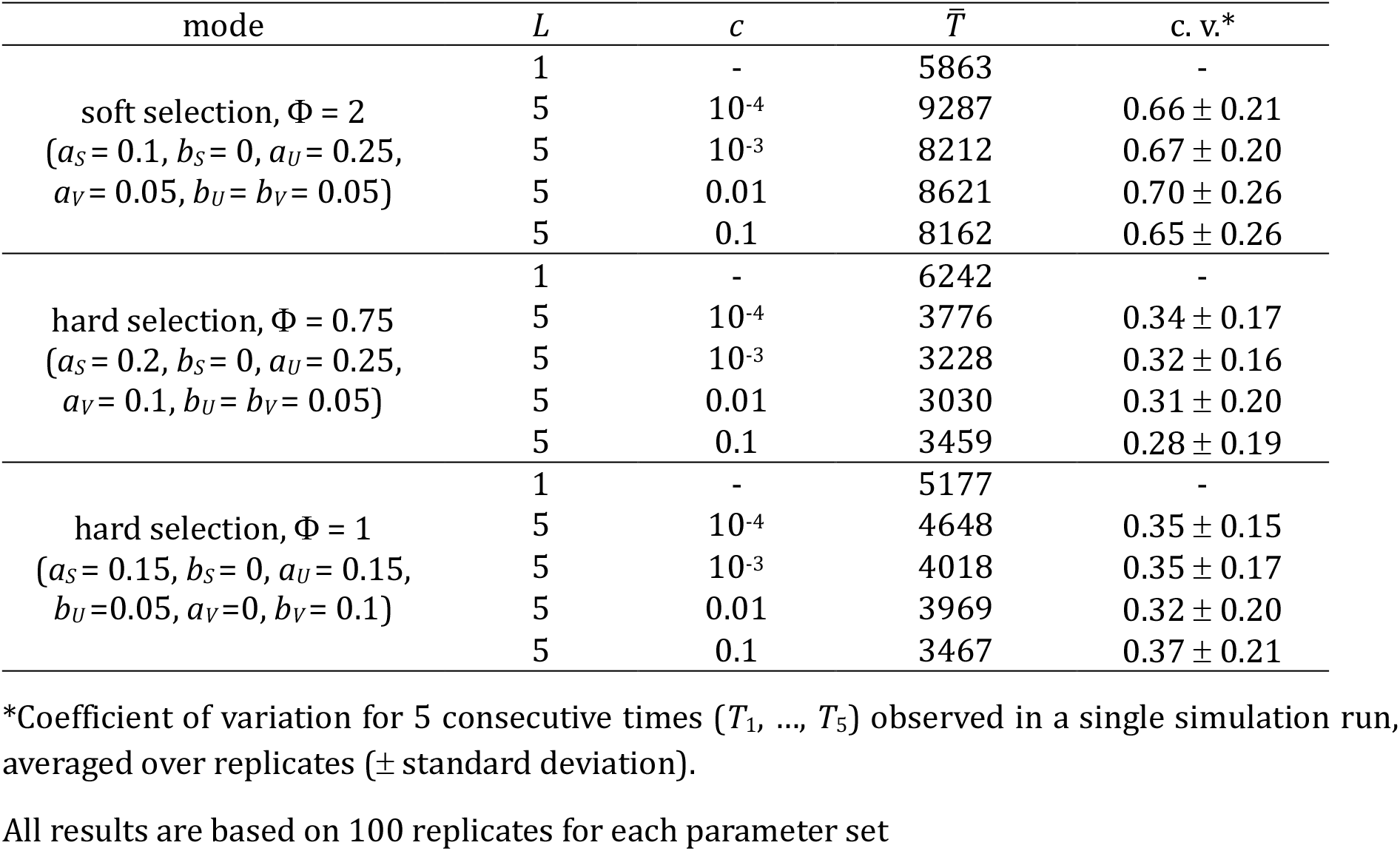
Mean time to the fixation (*q* > 0.9) of *A*_2_ in the multi-locus simulation.

**Table S2.**
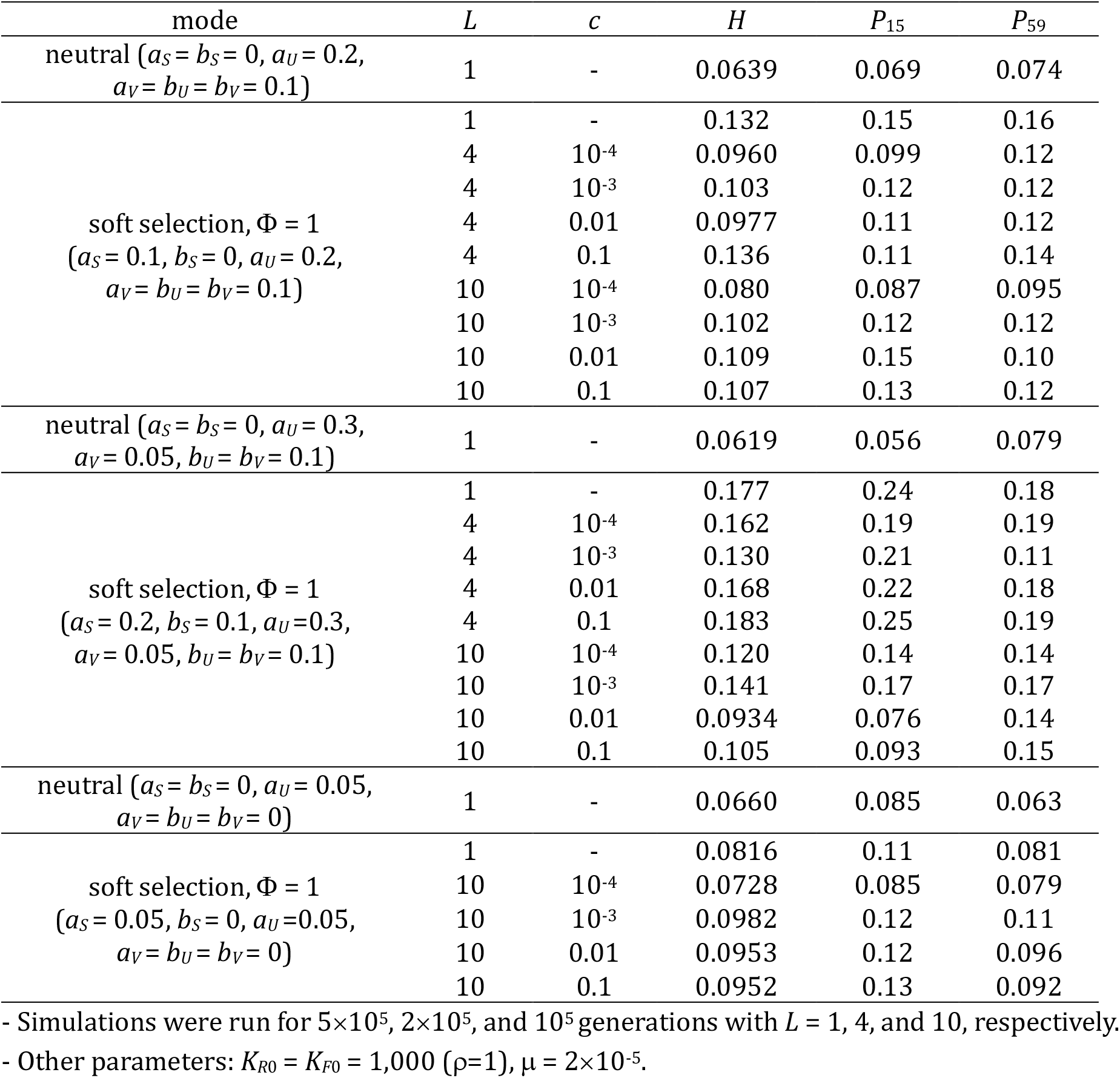
The long-term level of polymorphism in the multi-locus simulation.

**Figure S1:**
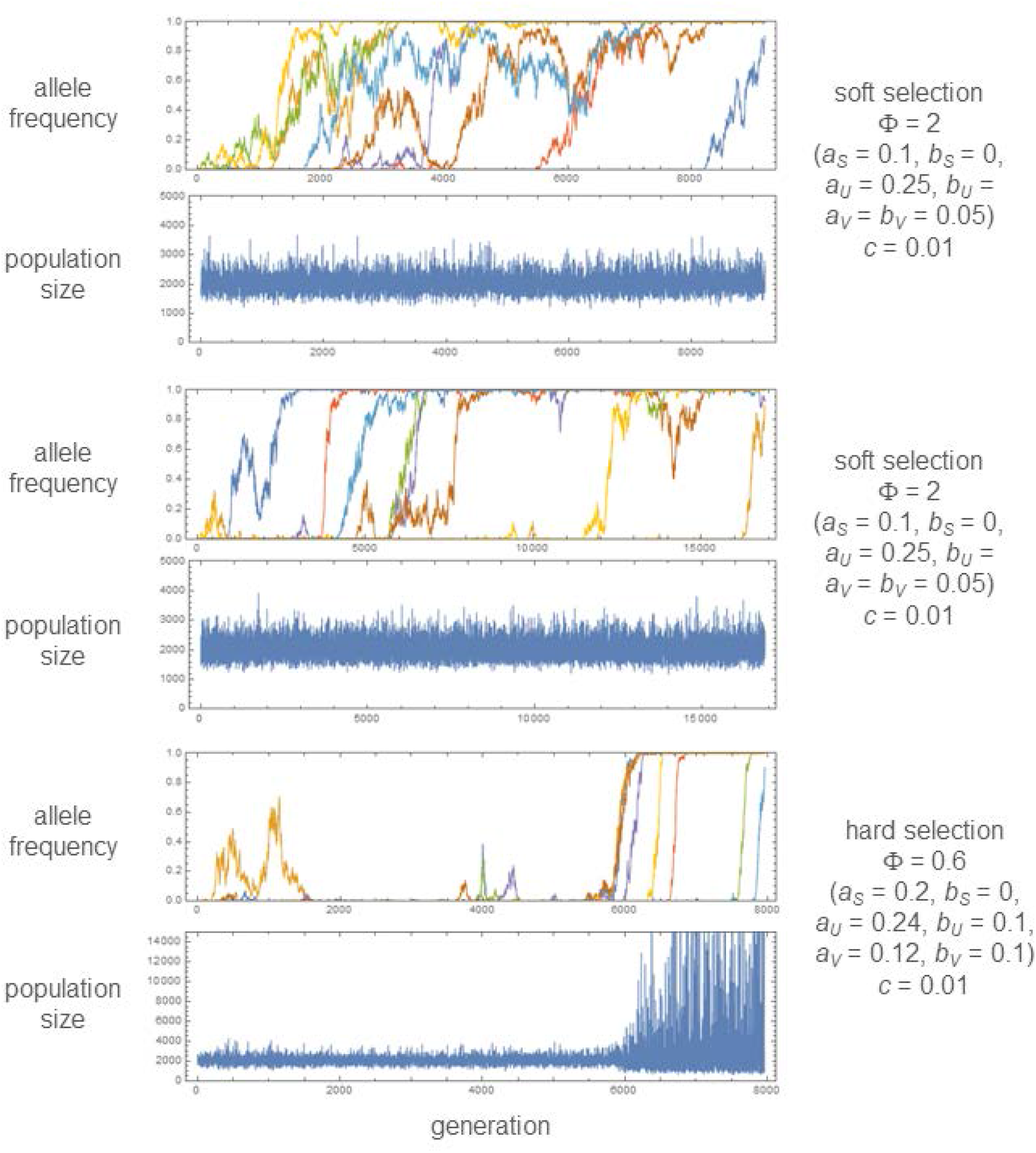
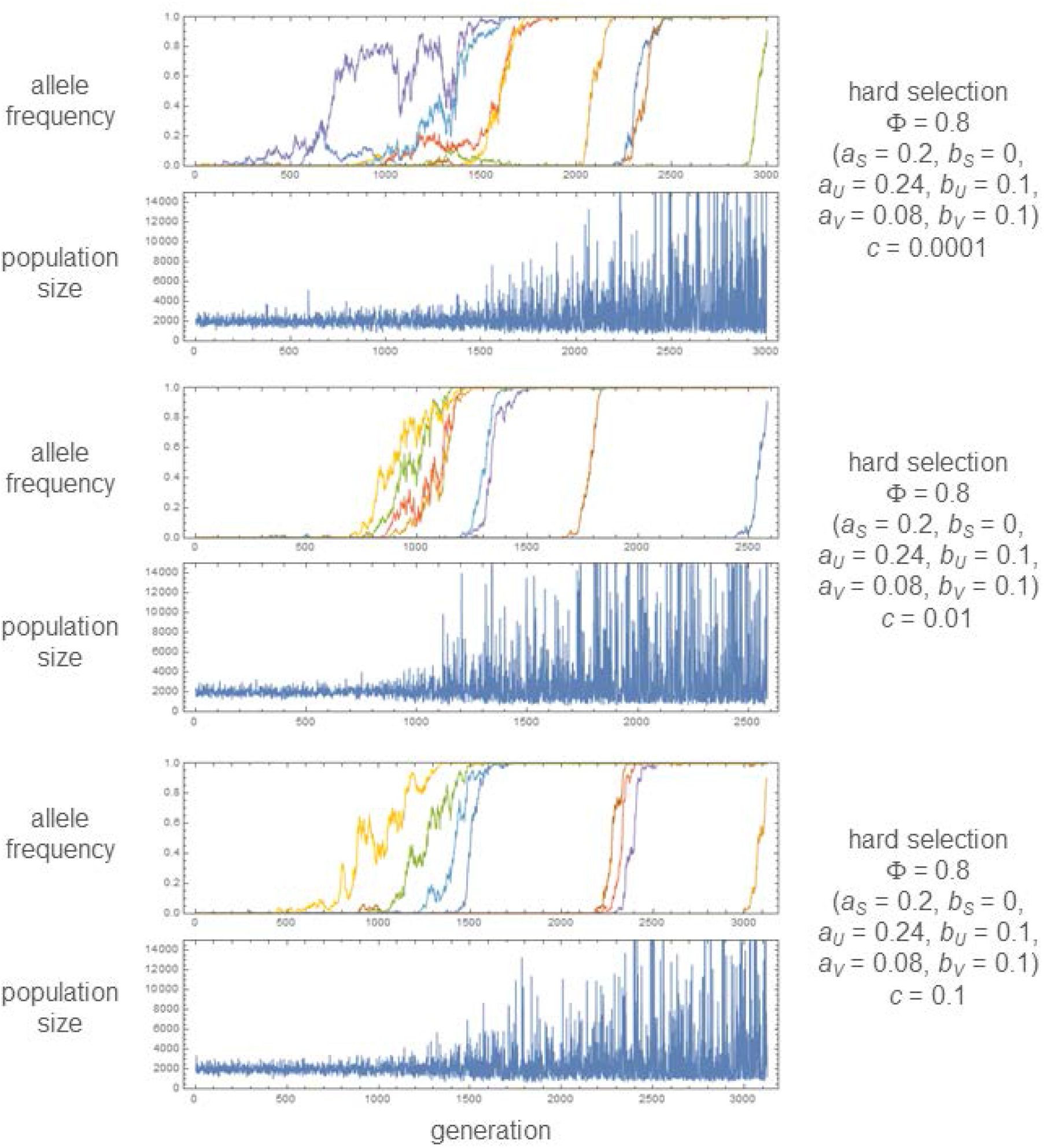
Examples of simulation runs in which the mutant allele (*A*_2_) reaches fixation sequentially at all 8 loci used. The change of population size is shown below that of allele frequencies, plotted in different colors for different loci. Parameter values are shown on the right side of graphs. Other parameters: *K*_*R*0_ = *K*_*F*0_ = 1000, μ = 2 × 10^−5^.

**Figure S2:**
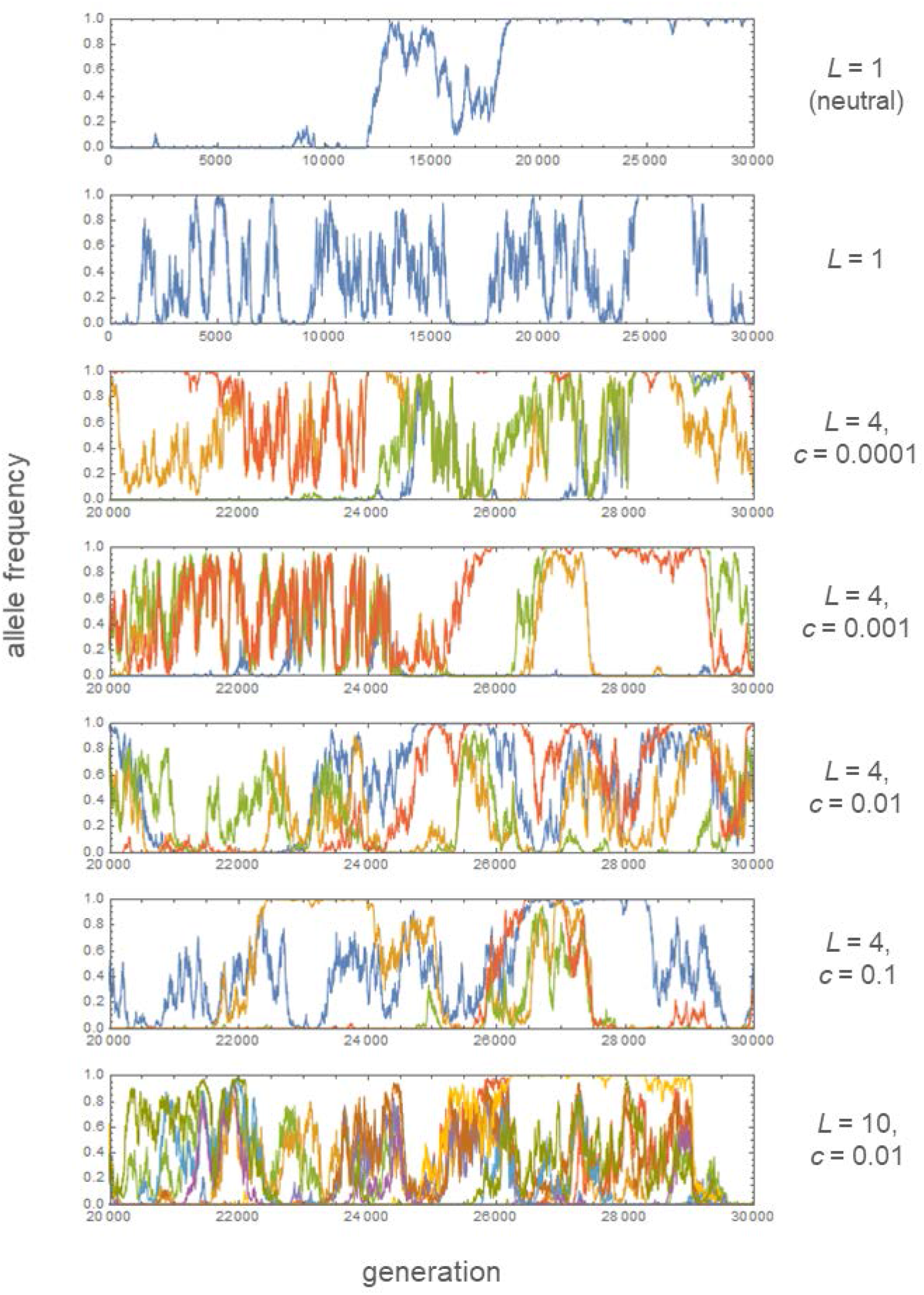
Exemplary trajectories of allele (*A*_2_) frequencies in simulations. Allele frequencies are plotted in different colors for different loci. The number of loci (*L*) and recombination rate (*c*) are shown on the right side of graphs. Other parameters: *K*_*R*0_ = *K*_*F*0_ = 1000, μ = 2×10^−5^, Φ = 1 (*a*_*S*_ = 0.2 [0 for neutral], *b*_*S*_ = 0.1, *a*_*U*_ = 0.3, *b*_*U*_ = 0.1, *a*_*V*_ = 0.05, *b*_*V*_ = 0.1).

## Notes

https://doi.org/10.5281/zenodo.12516522

### Competing Interest Statement

The authors have declared no competing interest.

### Summary of Updates

Several errors in referring equations were corrected

